## Supplemental methods and figures for "Transmission efficiency drives host-microbe associations"

***Authors for correspondence:**

#### Supplementary material:

#### 1. Supplementary methods

#### 2. Supplementary figures

#### 1. Supplementary methods

We consider here two discrete generation, deterministic mathematical models describing the evolution of microbe carrier and non-carrier frequencies in two well-mixed populations

(A and B). These models are defined to flexibly allow consideration of a range of different traits, that are detailed further below.

### (a) Mechanisms of microbe inheritance/uptake and transmission

Within these mathematical models it is assumed that there are two non-exclusive mechanisms by which individuals may become a microbe carrier – namely maternal transmission and environmental acquisition.

##### *Maternal transmission*

The first mechanism by which an individual may become a microbe carrier is via maternal transmission of the microbe. In this case, the offspring of a female microbe carrier will inherit the microbe with a probability ($\alpha$) in the range 0 to 1. For example, where $\alpha = 0.5$ the offspring of a microbe carrying mother will be half microbe carriers and half non-microbe carriers, regardless of the carrier status of the father. This maternal transmission probability is assumed to be equal for both male and female offspring. Note that in the case of asexual reproduction, the microbe can be vertically transmitted by all individuals and will be present in a fraction ($\alpha$) of their offspring.

##### *Environmental acquisition*

A second mechanism for an individual to become a microbe carrier is via environmental acquisition. Under this mechanism we assume that an individual acquires the microbe as a result of their actions within their environment – likely to be predominantly due to ingestion along with their diet. This effect is captured in the form of a probability of stable microbe uptake ($\beta$) in the range 0 to 1. This represents the probability that an individual will have acquired the microbe in the time period between their birth and their becoming sexually mature.

##### *Horizontal transmission*

The final mechanism by which an individual may become a microbe carrier in our models is via horizontal transmission. Here we assume that the microbe is acquired via contact between non-carrier and carrier individuals living in the same area. This is captured via the conversion of non-carrier individuals into carriers at a probability that varies depending upon the overall frequency of carrier individuals within the population. For example, a proportion $\tau(M_{1,A}^{t+1} + F_{1,A}^{t+1})$ of the non-carrier population will acquire the microbe as a result of horizontal transmission. Here $\tau$ is a parameter used to scale the strength of horizontal transmission by scaling the overall (i.e. both sexes) microbe carrier frequency into the appropriate probability of microbe acquisition. This process is assumed to occur following the application of maternal transmission, environmental acquisition and fitness costs but will act prior to any individuals dispersing and thus moving between populations.

### (b) Effects of the microbe on individual fitness

In addition to the three mechanisms of microbe uptake/acquisition detailed above, we consider the impact of microbe carrier status on host fitness. Within the model this is captured as a relative fitness ($\varepsilon$), representing a reduced survival resulting due to carrying the microbe. For instance, a relative fitness of $\varepsilon = 0.5$ means that microbe carrying individuals are half as likely to survive to sexual maturity than non-carrier individuals. Conversely, a relative fitness of $\varepsilon = 2$ represents a scenario in which microbe carrying individuals are twice as likely to survive to sexual maturity than non-carriers. Finally, a relative fitness of $\varepsilon = 1$ means that carrier and non-carrier individuals are equally likely to survive to sexual maturity.

### (c) Dispersal across populations

In the mathematical models considered here, we allow for bidirectional dispersion across two populations. This is captured as an exchange of some proportion ($\mu$) of the individuals in each population. For example, throughout this study we consider a value of $\mu = 0.02$ which means that 2% of the individuals from population A disperse into population B and simultaneously 2% of the individuals from population B disperse into population A. For simplicity we make several assumptions on the timing and form of this dispersal. Firstly, dispersal is assumed to occur once per generation, following the effects of microbe uptake/acquisition and fitness effects. We also assume that males and females are equally likely to disperse and that the two populations are well mixed such that the dispersing individuals will be a representative sample of each population. Finally, it is assumed that generations in each population are (approximately) synchronised such that all dispersal is simultaneous – meaning individuals may only move populations once within a single generation.

### (d) Modes of reproduction

Within this study we consider two distinct modes of reproduction. Specifically, we consider scenarios in which reproduction is exclusively asexual or exclusively sexual. In the case of sexual reproduction this necessitates the consideration of assortative mating, which would not be possible in an asexually reproducing population.

In both models we make a range of assumptions on the nature of the modelled populations. Firstly, it is assumed that each population is sufficiently large that stochastic effects and integer numbers of individuals may be neglected. These populations are then assumed to be panmictic (randomly mating) with a 1:1 (male to female) sex ratio both initially and in each subsequent generation. Due to the assumptions on the form of dispersal considered here, this 1:1 sex ratio is not able to be skewed by the effects of dispersal. Finally, it is assumed that population A consists of 1% microbe carriers and 99% non-microbe carriers whereas population B initially consists of exclusively non-microbe carriers.

##### *Asexual reproduction*

In the first instance we consider the simple case whereby all reproduction within our populations is exclusively asexual. This eliminates the requirement for us to consider male and female populations separately. As such, the model calculates the frequencies of non-microbe carrying individuals ($W_0$) and microbe carriers ($W_1$) in a two-step process. The first step is to calculate the relative proportions of the total population that will be of each type. These relative proportions are denoted by a superscript $e$ and are found using

$$W_{0,A}^{e} = \underbrace{(1-\beta_A)\, W_{0,A}^{t}}_{\substack{\text{Offspring of non-carrier not} \\ \text{acquiring from environment}}} + \underbrace{(1-\beta_A)(1-\alpha_A) W_{1,A}^{t}}_{\substack{\text{Offspring of carrier not acquiring} \\ \text{from environment and not inheriting} \\ \text{from parent}}} , \tag{1}$$

$$W_{1,A}^{e} = \varepsilon_A \left[ \underbrace{\beta_A W_{0,A}^{t}}_{\substack{\text{Offspring of non-carrier} \\ \text{acquiring from environment}}} + \underbrace{\beta_A (1-\alpha_A) W_{1,A}^{t}}_{\substack{\text{Offspring of carrier not inheriting} \\ \text{from parent but acquiring from} \\ \text{environment}}} + \underbrace{\alpha_A W_{1,A}^{t}}_{\substack{\text{Offspring of carrier} \\ \text{inheriting from parent}}} \right], \tag{2}$$

where each of the parameter symbols are as described in the sections above and the superscript $t$ denotes the time point used for calculations (i.e. the previous generation). Note that these equations are for population A (hence the subscript $A$) and there is also a pair of equations for population B that are identical to equations (1) and (2) except that they have the $A$ subscript with a $B$ subscript.

Step two in this process is to normalise these relative proportions such that they fill the range zero to one. Thus, the frequencies of microbe carriers and non-carriers are obtained using

$$\widetilde{W}_{0,A}^{t+1} = \frac{W_{0,A}^{e}}{\Omega_A}, \quad (3)$$

$$\widetilde{W}_{1,A}^{t+1} = \frac{W_{1,A}^{e}}{\Omega_A}, \quad (4)$$

$$\widetilde{W}_{0,B}^{t+1} = \frac{W_{0,B}^{e}}{\Omega_B}, \quad (5)$$

$$\widetilde{W}_{1,B}^{t+1} = \frac{W_{0,B}^{e}}{\Omega_B}, \quad (6)$$

where $\Omega_A$ and $\Omega_B$ represent the overall fitness of each population and are given by

$$\Omega_A = W_{0,A}^{e} + W_{1,A}^{e}, \quad (7)$$

$$\Omega_B = W_{0,B}^{e} + W_{1,B}^{e}. \quad (8)$$

These normalised frequencies are then used to calculate the amount of horizontal transmission that occurs within the population. This is achieved using

$$\underline{W}_{0,A}^{t+1} = \left(1 - \tau\, \widetilde{W}_{1,A}^{t+1}\right) \widetilde{W}_{0,A}^{t+1}, \quad (9)$$

$$\underline{W}_{1,A}^{t+1} = \widetilde{W}_{1,A}^{t+1} + \tau\, \widetilde{W}_{1,A}^{t+1} \widetilde{W}_{0,A}^{t+1}, \quad (10)$$

$$\underline{W}_{0,B}^{t+1} = \left(1 - \tau\, \widetilde{W}_{1,B}^{t+1}\right) \widetilde{W}_{0,B}^{t+1}, \quad (11)$$

$$\underline{W}_{1,B}^{t+1} = \widetilde{W}_{1,B}^{t+1} + \tau\, \widetilde{W}_{1,B}^{t+1} \widetilde{W}_{0,B}^{t+1}. \quad (12)$$

We then obtain our final frequencies for a given generation by performing the dispersal calculations according to

$$W_{0,A}^{t+1} = (1-\mu)\underline{W}_{0,A}^{t+1} + \mu \underline{W}_{0,B}^{t+1}, \quad (13)$$

$$W_{1,A}^{t+1} = (1-\mu)\underline{W}_{1,A}^{t+1} + \mu \underline{W}_{1,B}^{t+1}, \quad (14)$$

$$W_{0,B}^{t+1} = (1-\mu)\underline{W}_{0,B}^{t+1} + \mu \underline{W}_{0,A}^{t+1}, \quad (15)$$

$$W_{0,B}^{t+1} = (1-\mu)\underline{W}_{0,B}^{t+1} + \mu \underline{W}_{0,A}^{t+1}. \quad (16)$$

Within the above expressions the ~ and – symbols over the $W$ variables are used to signify temporary variables that are simply used as intermediates used in moving from one model stage to the next.

Note that we consider fitness effects and rates of vertical transmission to be intrinsic properties of the microbe and the species in question rather than the environment, meaning that $\varepsilon_A = \varepsilon_B$ and $\alpha_A = \alpha_B$.

#### *Sexual reproduction*

We then go on to consider the case in which all reproduction within the two populations is exclusively sexual. Unlike the asexual example above, here we must consider male and female populations separately. As such, this model calculates the frequencies of non-microbe carrying males ($M_0$) and females ($F_0$) as well as microbe carrying males ($M_1$) and females ($F_1$) in a two-step process. The first step is to calculate the relative proportions of the total population that will be of each type. As in the asexual reproduction case, these relative proportions are denoted by a subscript $e$ and are calculated using

$$M_{0,A}^{e} = \frac{1}{2}\left[\underbrace{M_{0,A}^{t}(1-\beta_A)P_{0,0}}_{\substack{\text{Offspring of non-carrier}\\ \text{mother not acquiring from}\\ \text{the environment}}} + \underbrace{M_{0,A}^{t}(1-\alpha_A)(1-\beta_A)P_{0,1}}_{\substack{\text{Offspring of carrier mother not}\\ \text{acquiring from the environment}\\ \text{and not inheriting from mother}}} + \underbrace{M_{1,A}^{t}(1-\beta_A)P_{1,0}}_{\substack{\text{Offspring of non-carrier}\\ \text{mother not acquiring}\\ \text{from the environment}}} + \underbrace{M_{1,A}^{t}(1-\alpha_A)(1-\beta_A)P_{1,1}}_{\substack{\text{Offspring of carrier mother not}\\ \text{acquiring from the environment}\\ \text{and not inheriting from mother}}}\right], \tag{17}$$

$$M_{1,A}^{e} = \frac{1}{2}\varepsilon_A\left[\begin{array}{c}\underbrace{M_{0,A}^{t}\beta_A P_{0,0}}_{\substack{\text{Offspring of non-carrier}\\ \text{mother acquiring from}\\ \text{the environment}}} + \underbrace{M_{0,A}^{t}\alpha_A P_{0,1}}_{\substack{\text{Offspring of carrier}\\ \text{mother inheriting}\\ \text{from mother}}} + \underbrace{M_{0,A}^{t}(1-\alpha_A)\beta_A P_{0,1}}_{\substack{\text{Offspring of carrier mother}\\ \text{not inheriting but acquiring}\\ \text{from the environment}}} + \underbrace{M_{1,A}^{t}\beta_A P_{1,0}}_{\substack{\text{Offspring of non-carrier}\\ \text{mother acquiring from}\\ \text{the environment}}}\\ + \underbrace{M_{1,A}^{t}\alpha_A P_{1,1}}_{\substack{\text{Offspring of carrier}\\ \text{mother inheriting}\\ \text{from mother}}} + \underbrace{M_{1,A}^{t}(1-\alpha_A)\beta_A P_{1,1}}_{\substack{\text{Offspring of carrier mother}\\ \text{not inheriting from mother}\\ \text{but acquiring from environment}}}\end{array}\right], \tag{18}$$

where all parameter symbols are as described in the previous sections and the superscript $t$ denotes the time point used for calculations (i.e. the previous generation). Note that these equations are for males in population A (hence the subscript $A$). There is also an identical pair of equations for females in population A and a set of four equations for population B in which the $A$ subscript in the above is replaced with a $B$. Within these equations the symbols $P_{i,j}$ are used to represent the probability of mating with a certain type (i.e. carrier or non-carrier) under a given regime of assortative mating – with full details given below.

Step two in this process is to normalise the relative proportions such that they fill the range zero to one. Thus, the frequencies of microbe carrier and non-carrier males and females are obtained using

$$\widetilde{M}_{0,A}^{t+1} = \frac{M_{0,A}^{e}}{\Omega_A}, \tag{19}$$

$$\widetilde{M}_{1,A}^{t+1} = \frac{M_{1,A}^{e}}{\Omega_A}, \tag{20}$$

$$\widetilde{F}_{0,A}^{t+1} = \frac{F_{0,A}^{e}}{\Omega_A}, \tag{21}$$

$$\widetilde{F}_{1,A}^{t+1} = \frac{F_{1,A}^{e}}{\Omega_A}, \tag{22}$$

$$\widetilde{M}_{0,B}^{t+1} = \frac{M_{0,B}^{e}}{\Omega_B}, \tag{23}$$

$$\widetilde{M}_{1,B}^{t+1} = \frac{M_{1,B}^{e}}{\Omega_B}, \tag{24}$$

$$\widetilde{F}_{0,B}^{t+1} = \frac{F_{0,B}^{e}}{\Omega_B}, \tag{25}$$

$$\widetilde{F}_{1,B}^{t+1} = \frac{F_{1,B}^{e}}{\Omega_B}, \tag{26}$$

where $\Omega_A$ and $\Omega_B$ represent the overall fitness of each population and are given by

$$\Omega_A = M_{0,A}^{e} + M_{1,A}^{e} + F_{0,A}^{e} + F_{1,A}^{e}, \tag{27}$$

$$\Omega_B = M_{0,B}^{e} + M_{1,B}^{e} + F_{0,B}^{e} + F_{1,B}^{e}. \tag{28}$$

These normalised frequencies are then used to calculate the effects of horizontal transmission within the modelled populations. This is achieved using

$$\underline{M}_{0,A}^{t+1} = \left(1 - \tau\left(\widetilde{M}_{1,A}^{t+1} + \widetilde{F}_{1,A}^{t+1}\right)\right)\widetilde{M}_{0,A}^{t+1}, \tag{29}$$

$$\underline{M}_{1,A}^{t+1} = \widetilde{M}_{1,A}^{t+1} + \tau\left(\widetilde{M}_{1,A}^{t+1} + \widetilde{F}_{1,A}^{t+1}\right)\widetilde{M}_{0,A}^{t+1}, \tag{30}$$

$$\underline{F}_{0,A}^{t+1} = \left(1 - \tau\left(\widetilde{M}_{1,A}^{t+1} + \widetilde{F}_{1,A}^{t+1}\right)\right)\widetilde{F}_{0,A}^{t+1}, \tag{31}$$

$$\underline{F}_{1,A}^{t+1} = \widetilde{F}_{1,A}^{t+1} + \tau\left(\widetilde{M}_{1,A}^{t+1} + \widetilde{F}_{1,A}^{t+1}\right)\widetilde{F}_{0,A}^{t+1}, \tag{32}$$

$$\underline{M}_{0,B}^{t+1} = \left(1 - \tau\left(\widetilde{M}_{1,B}^{t+1} + \widetilde{F}_{1,B}^{t+1}\right)\right)\widetilde{M}_{0,B}^{t+1}, \tag{33}$$

$$\underline{M}_{1,B}^{t+1} = \widetilde{M}_{1,B}^{t+1} + \tau\left(\widetilde{M}_{1,B}^{t+1} + \tilde{F}_{1,B}^{t+1}\right)\widetilde{M}_{0,B}^{t+1}, \tag{34}$$
$$\underline{F}_{0,B}^{t+1} = \left(1 - \tau\left(\widetilde{M}_{1,B}^{t+1} + \tilde{F}_{1,B}^{t+1}\right)\right)\tilde{F}_{0,B}^{t+1}, \tag{35}$$
$$\underline{F}_{1,B}^{t+1} = \tilde{F}_{1,B}^{t+1} + \tau\left(\widetilde{M}_{1,B}^{t+1} + \tilde{F}_{1,B}^{t+1}\right)\tilde{F}_{0,B}^{t+1}. \tag{36}$$

We then obtain our final frequencies by performing dispersal calculations according to

$$M_{0,A}^{t+1} = (1-\mu)\underline{M}_{0,A}^{t+1} + \mu\underline{M}_{0,B}^{t+1}, \tag{37}$$
$$M_{1,A}^{t+1} = (1-\mu)\underline{M}_{1,A}^{t+1} + \mu\underline{M}_{1,B}^{t+1}, \tag{38}$$
$$F_{0,A}^{t+1} = (1-\mu)\underline{F}_{0,A}^{t+1} + \mu\underline{F}_{0,B}^{t+1}, \tag{39}$$
$$F_{1,A}^{t+1} = (1-\mu)\underline{F}_{1,A}^{t+1} + \mu\underline{F}_{1,B}^{t+1}, \tag{40}$$
$$M_{0,B}^{t+1} = (1-\mu)\underline{M}_{0,B}^{t+1} + \mu\underline{M}_{0,A}^{t+1}, \tag{41}$$
$$M_{1,B}^{t+1} = (1-\mu)\underline{M}_{1,B}^{t+1} + \mu\underline{M}_{1,A}^{t+1}, \tag{42}$$
$$F_{0,B}^{t+1} = (1-\mu)\underline{F}_{0,B}^{t+1} + \mu\underline{F}_{0,A}^{t+1}, \tag{43}$$
$$F_{1,B}^{t+1} = (1-\mu)\underline{F}_{1,B}^{t+1} + \mu\underline{F}_{1,A}^{t+1}, \tag{44}$$

Within the above expressions the ~ and – symbols over the $W$ variables are used to signify temporary variables that are simply used as intermediates used in moving from one model stage to the next.

Note that we consider fitness effects and rates of vertical transmission to be intrinsic properties of the microbe and the species in question rather than the environment, meaning that $\varepsilon_A = \varepsilon_B$ and $\alpha_A = \alpha_B$.

### (e) Host mate choice - assortative mating

To capture the effects of assortative mating within the mathematical model of a sexually reproducing population we consider an n-choice mating framework similar to that of Newberry et al. (2016). Within this framework an individual is given $n$ chances to find an individual of their preferred type to mate with. As the equations in the above section are formulated with the choice of mate going to the males, we first consider a scenario whereby males are given $n$ chances to find a microbe carrying female. Here, the male will encounter $n$ females sequentially and will mate if any of the first $n-1$ females sampled is a microbe carrier. On encountering their final (i.e. $n$-th) sampled female, the male will mate regardless of the microbe carrying status of that female. This essentially represents a fixed maximum energy investment that an individual is willing to make in order to find a mate of their preferred type.

In total we consider four distinct forms of assortative mating – each of which follows the same general framework as described in the example above. The first is that described in the above example whereby males prefer to mate with female microbe carriers and are given $n$ chances to do so. Mathematically this can be described as

$$P_{i,j} = F_{0,k}^{t}\left(1 - F_{1,k}^{t}\right)^{n-1}, when\ j = 0\ and$$
$$P_{i,j} = 1 - \left(1 - F_{1,k}^{t}\right)^{n}, when\ j = 1,$$

where $k$ (= $A\ or\ B$) denoting the population in which the assortative mating is occurring.

The other scenario in which males can sample $n$ females to find their preferred mating type is that whereby they prefer to mate with an individual of the same carrier status (i.e. (non-)carriers prefer to mate with (non-)carriers). This gives expressions for $P_{i,j}$ of the form

$$P_{i,j} = 1 - \left(1 - F_{j,k}^{t}\right)^{n}, when\ i = j\ and$$
$$P_{i,j} = F_{j,k}^{t}\left(1 - F_{i,k}^{t}\right)^{n-1}, when\ i \neq j,$$

where again $k$ (= $A\ or\ B$) denotes the population in which the assortative mating is occurring.

In addition to the male-choice scenarios considered above, we also consider the same two preferences but, in the case whereby females can sample $n$ males in order to find a mate of their preferred type. Mathematically, this leads to the same expressions listed above but with the $F$ symbols swapped for $M$.

### 2. Supplementary figures

**Fig S1A. Effect of transmission modes and microbe effects on host fitness on host-microbe carrier frequencies within a single population of asexually reproducing hosts.** Here each line represents different steady-state microbe carrier frequencies (0.1-0.9, increasing in 0.2 intervals). From top to bottom: increasing levels of horizontal transmission for 0-0.5. From left to right: relative fitness of carriers to non-carriers 0.5-2. Effects of asexual reproduction (compare to Figs S2A, S3A, S4A & S5A) on carrier frequencies are modest.

**Fig S1B. Effect of transmission modes and microbe effects on host fitness on host-microbe carrier frequencies within two populations of asexually reproducing hosts.** Here each line represents different steady-state microbe carrier frequencies (0.1-0.9, increasing in 0.2 intervals). From top to bottom: increasing levels of horizontal transmission for 0-0.5. From left to right: relative fitness of carriers to non-carriers 0.5-2. In population 1 microbes can proliferate independently of hosts and there is environmental acquisition by the host. In population 2 microbes cannot be acquired by hosts from the environment, only via maternal VT or horizontal transmission from migrants from population 1. Effects of asexual reproduction (compare to Figs S2B, S3B, S4B & S5B) on carrier frequencies are modest.

**Fig S2A. Effect of transmission modes, host mate choice, and microbe effects on host fitness on host-microbe carrier frequencies within a single population of sexually reproducing hosts. Females are the choosing sex and prefer males that are microbe-carriers.** Here each line represents different steady-state microbe carrier frequencies (0.1-0.9, increasing in 0.2 intervals). From top to bottom: increasing levels of horizontal transmission for 0-0.5. From left to right: relative fitness of carriers to non-carriers 0.5-2. Here mate choice produces no effect on carrier frequencies under set rates of acquisition, as females are the sex producing VT and choice of partner (carrier or non-carrier) does not alter transmission (compare to Fig S4A).

**Fig S2B. Effect of transmission modes, host mate choice, and microbe effects on host fitness on host-microbe carrier frequencies within two populations of sexually reproducing hosts. Females are the choosing sex and prefer males that are microbe-carriers.** Here each line represents different steady-state microbe carrier frequencies (0.1-0.9, increasing in 0.2 intervals). From top to bottom: increasing levels of horizontal transmission for 0-0.5. From left to right: relative fitness of carriers to non-carriers 0.5-2. In population 1 microbes can proliferate independently of hosts and there is environmental acquisition by the host. In population 2 microbes cannot be acquired by hosts from the environment, only via maternal VT or horizontal transmission from migrants from population 1.

**Fig S3A. Effect of transmission modes, host mate choice, and microbe effects on host fitness on host-microbe carrier frequencies within a single population of sexually reproducing hosts. Females are the choosing sex and prefer males of the same carrier/non-carrier status.** Here each line represents different steady-state microbe carrier frequencies (0.1-0.9, increasing in 0.2 intervals). From top to bottom: increasing levels of horizontal transmission for 0-0.5. From left to right: relative fitness of carriers to non-carriers 0.5-2. Here the effects of mate choice on carrier frequencies are less pronounced than for male mate choice compare with Fig S5A.

**Fig S3B. Effect of transmission modes, host mate choice, and microbe effects on host fitness on host-microbe carrier frequencies within two populations of sexually reproducing hosts. Females are the choosing sex and prefer males of the same carrier/non-carrier status.** Here each line represents different steady-state microbe carrier frequencies (0.1-0.9, increasing in 0.2 intervals). From top to bottom: increasing levels of horizontal transmission for 0-0.5. From left to right: relative fitness of carriers to non-carriers 0.5-2. In population 1 microbes can proliferate independently of hosts and there is environmental acquisition by the host. In population 2 microbes cannot be acquired by hosts from the environment, only via maternal VT or horizontal transmission from migrants from population 1.

**Fig S4A. Effect of transmission modes, host mate choice, and microbe effects on host fitness, on host-microbe carrier frequencies within a single population of sexually reproducing hosts. Males are the choosing sex and prefer females that are microbe-carriers.** Here each line represents different steady-state microbe carrier frequencies (0.1-0.9, increasing in 0.2 intervals). From top to bottom: increasing levels of horizontal transmission for 0-0.5. From left to right: relative fitness of carriers to non-carriers 0.5-2. Top row panels 2 & 4 (Left to Right) are reproduced in main text Fig 2C and D.

**Fig S4B. Effect of transmission modes, host mate choice, and microbe effects on host fitness on host-microbe carrier frequencies within two populations of sexually reproducing hosts. Males are the**

**choosing sex and prefer females that are microbe-carriers.** Here each line represents different steady-state microbe carrier frequencies (0.1-0.9, increasing in 0.2 intervals). From top to bottom: increasing levels of horizontal transmission for 0-0.5. From left to right: relative fitness of carriers to non-carriers 0.5-2. In population 1 microbes can proliferate independently of hosts and there is environmental acquisition by the host. In population 2 microbes cannot be acquired by hosts from the environment, only via maternal VT or horizontal transmission from migrants from population 1. Top row panels 3 & 5 (Left to Right) are reproduced in main text Fig 3C and D.

**Fig S5A. Effect of transmission modes, host mate choice, and microbe effects on host fitness on host-microbe carrier frequencies within a single population of sexually reproducing hosts. Males are the choosing sex and prefer females of the same carrier/non-carrier status.** Here each line represents different steady-state microbe carrier frequencies (0.1-0.9, increasing in 0.2 intervals). From top to bottom: increasing levels of horizontal transmission for 0-0.5. From left to right: relative fitness of carriers to non-carriers 0.5-2. Top row panels 2 & 4 (Left to Right) are reproduced in main text Fig 2A and B.

**Fig S5B. Effect of transmission modes, host mate choice, and microbe effects on host fitness on host-microbe carrier frequencies within two populations of sexually reproducing hosts. Males are the choosing sex and prefer females of the same carrier/non-carrier status.** Here each line represents different steady-state microbe carrier frequencies (0.1-0.9, increasing in 0.2 intervals). From top to bottom: increasing levels of horizontal transmission for 0-0.5. From left to right: relative fitness of carriers to non-carriers 0.5-2. In population 1 microbes can proliferate independently of hosts and there is environmental acquisition by the host. In population 2 microbes cannot be acquired by hosts from the environment, only via maternal VT or horizontal transmission from migrants from population 1. Top row panels 3 & 5 (Left to Right) are reproduced in main text Fig 3A and B.

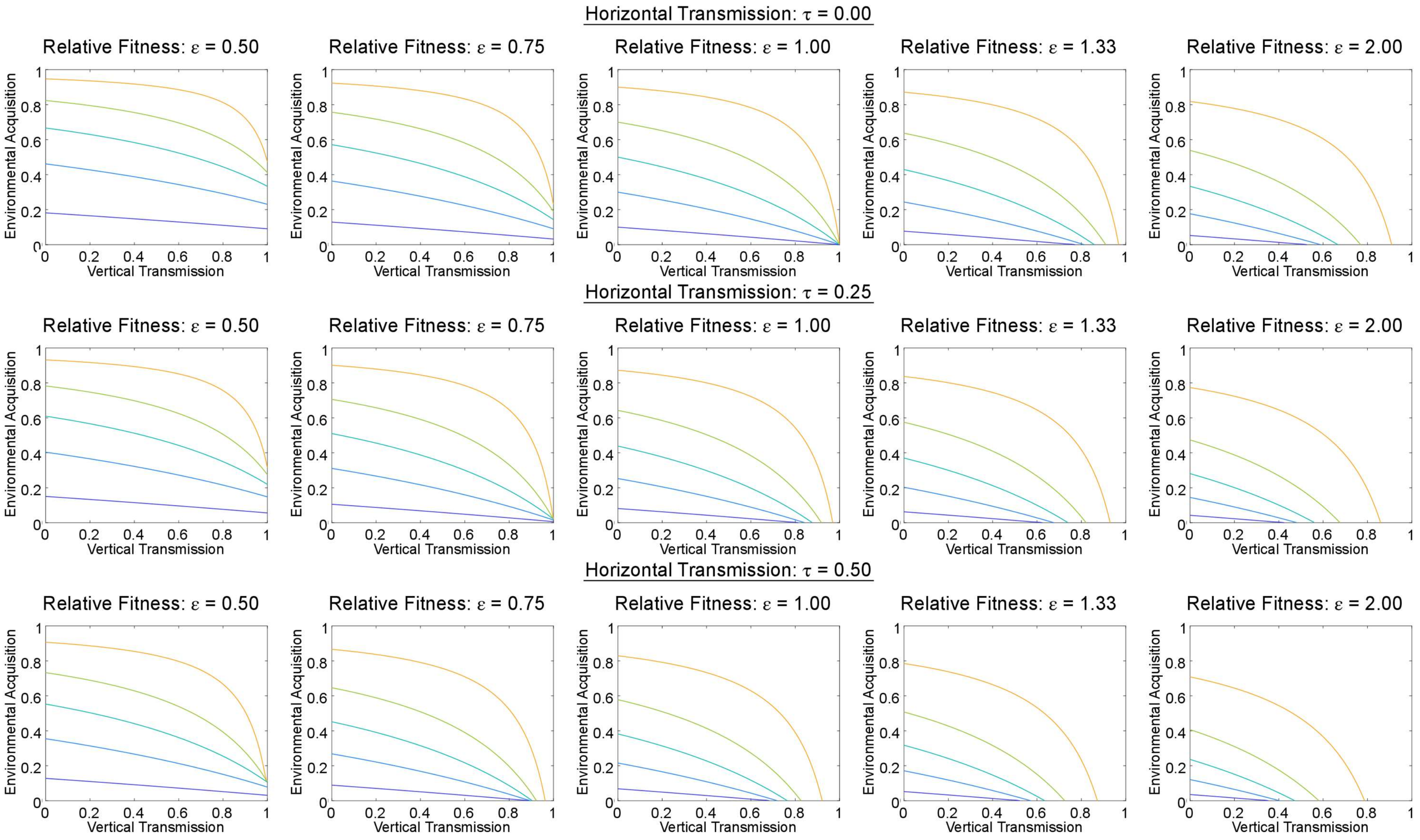


**Fig S1A. Effect of transmission modes and microbe effects on host fitness on host-microbe carrier frequencies within a single population of asexually reproducing hosts.** Here each line represents different steady-state microbe carrier frequencies (0.1-0.9, increasing in 0.2 intervals). From top to bottom: increasing levels of horizontal transmission for 0-0.5. From left to right: relative fitness of carriers to non-carriers 0.5-2. Effects of asexual reproduction (compare to Figs S2, 4, 6 & 8) on carrier frequencies are modest.

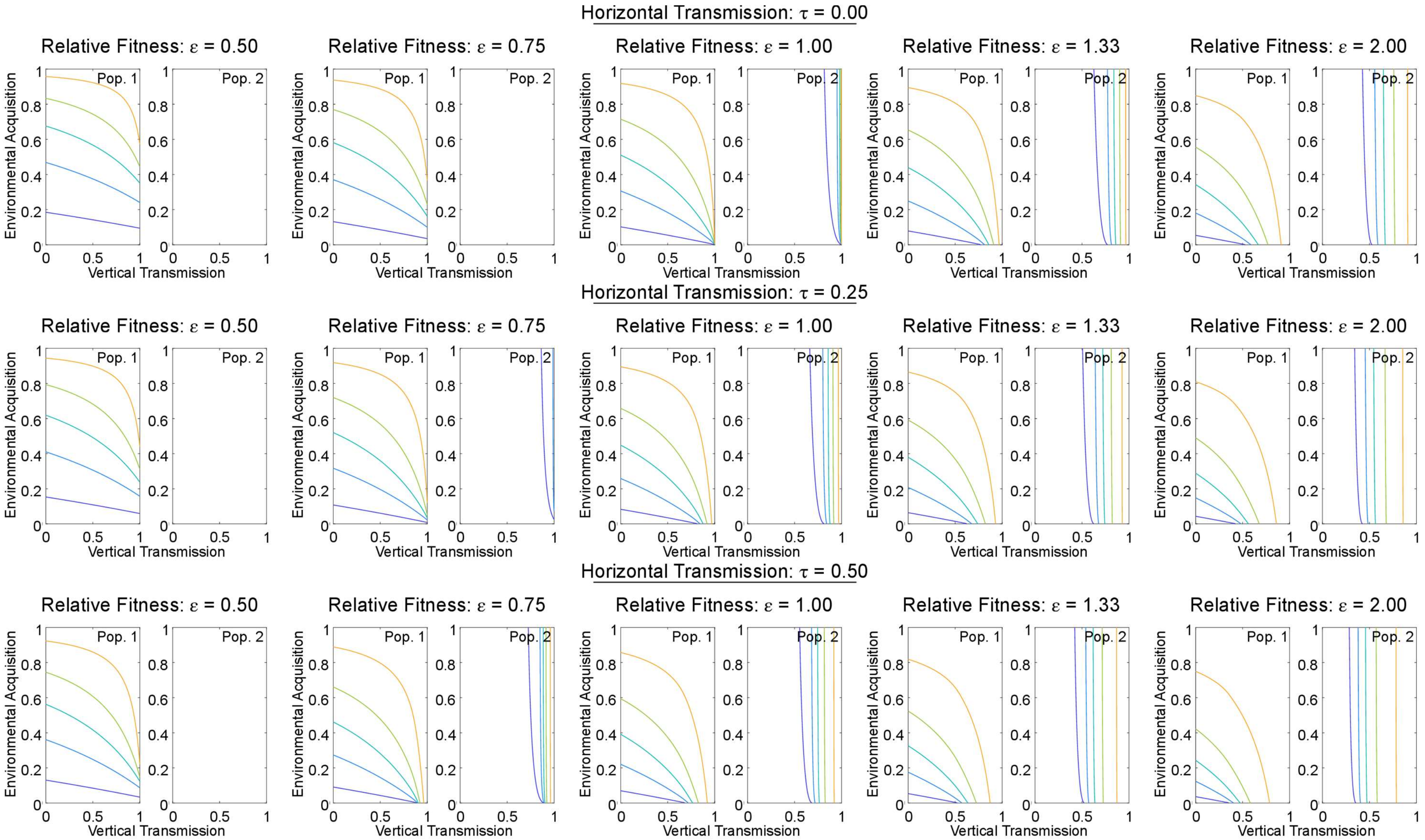


**Fig S1B. Effect of transmission modes and microbe effects on host fitness on host-microbe carrier frequencies within two populations of asexually reproducing hosts.** Here each line represents different steady-state microbe carrier frequencies (0.1-0.9, increasing in 0.2 intervals). From top to bottom: increasing levels of horizontal transmission for 0-0.5. From left to right: relative fitness of carriers to non-carriers 0.5-2. In population 1 microbes can proliferate independently of hosts and there is environmental acquisition by the host. In population 2 microbes cannot be acquired by hosts from the environment, only via maternal VT or horizontal transmission from migrants from population 1. Effects of asexual reproduction (compare to Figs S3, 5, 7 & 9) on carrier frequencies are modest.

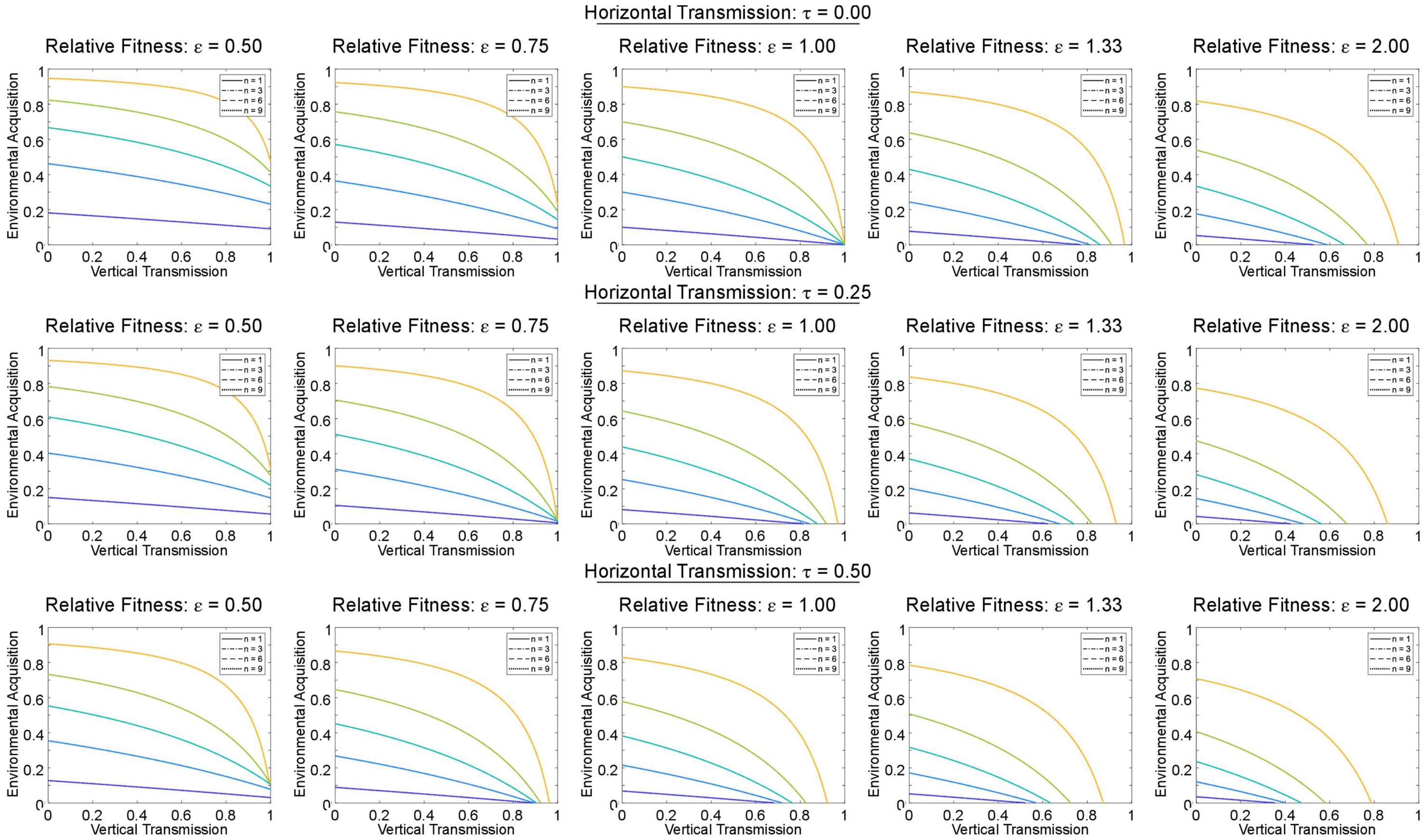


**Fig S2A. Effect of transmission modes, host mate choice, and microbe effects on host fitness on host-microbe carrier frequencies within a single population of sexually reproducing hosts. Females are the choosing sex and prefer males that are microbe-carriers.** Here each line represents different steady-state microbe carrier frequencies (0.1-0.9, increasing in 0.2 intervals). From top to bottom: increasing levels of horizontal transmission for 0-0.5. From left to right: relative fitness of carriers to non-carriers 0.5-2. Here mate choice produces no effect on carrier frequencies under set rates of acquisition, as females are the sex producing VT and choice of partner (carrier or non-carrier) does not alter transmission (compare to Fig S6).

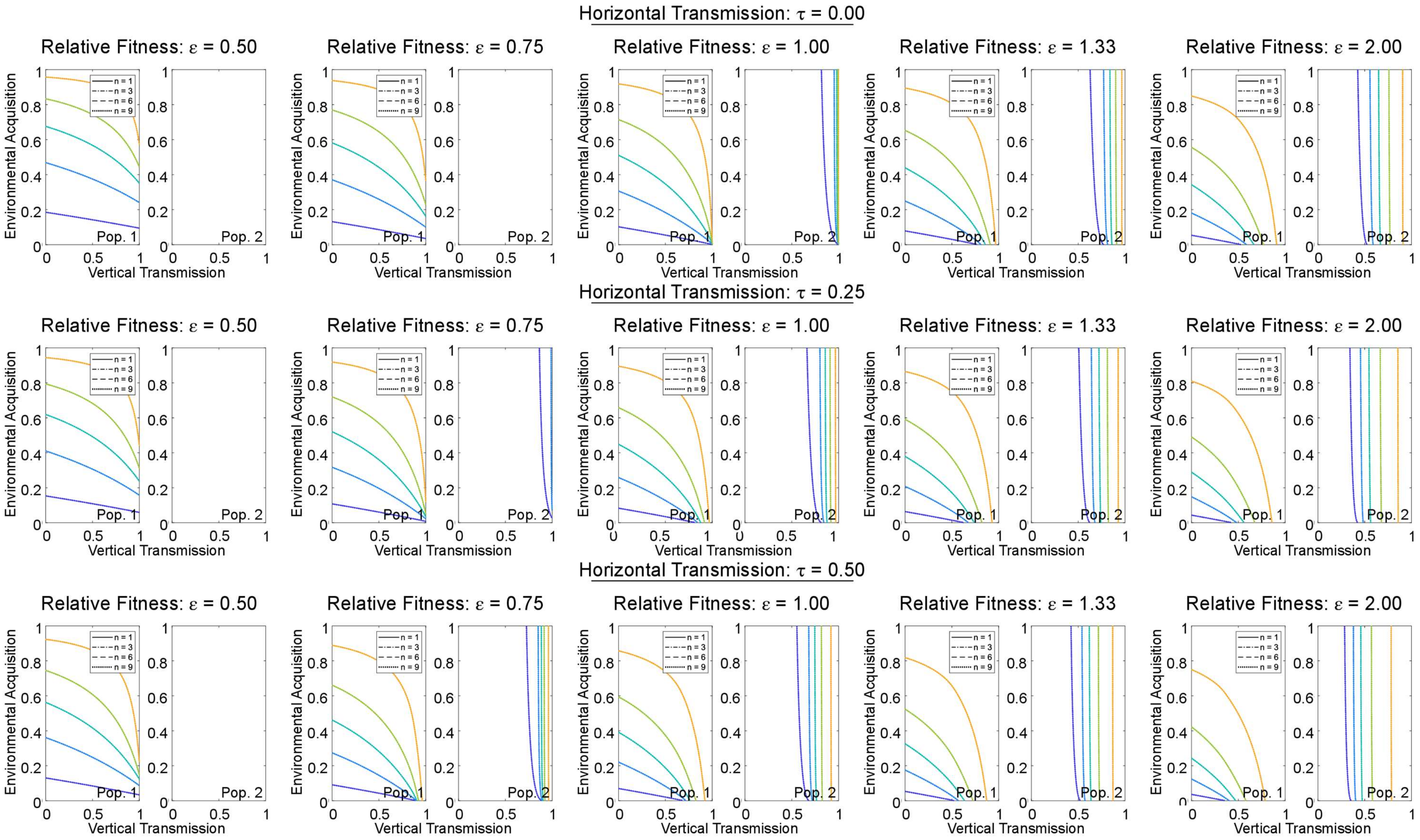


**Fig S2B. Effect of transmission modes, host mate choice, and microbe effects on host fitness on host-microbe carrier frequencies within two populations of sexually reproducing hosts. Females are the choosing sex and prefer males that are microbe-carriers.** Here each line represents different steady-state microbe carrier frequencies (0.1-0.9, increasing in 0.2 intervals). From top to bottom: increasing levels of horizontal transmission for 0-0.5. From left to right: relative fitness of carriers to non-carriers 0.5-2. In population 1 microbes can proliferate independently of hosts and there is environmental acquisition by the host. In population 2 microbes cannot be acquired by hosts from the environment, only via maternal VT or horizontal transmission from migrants from population 1.

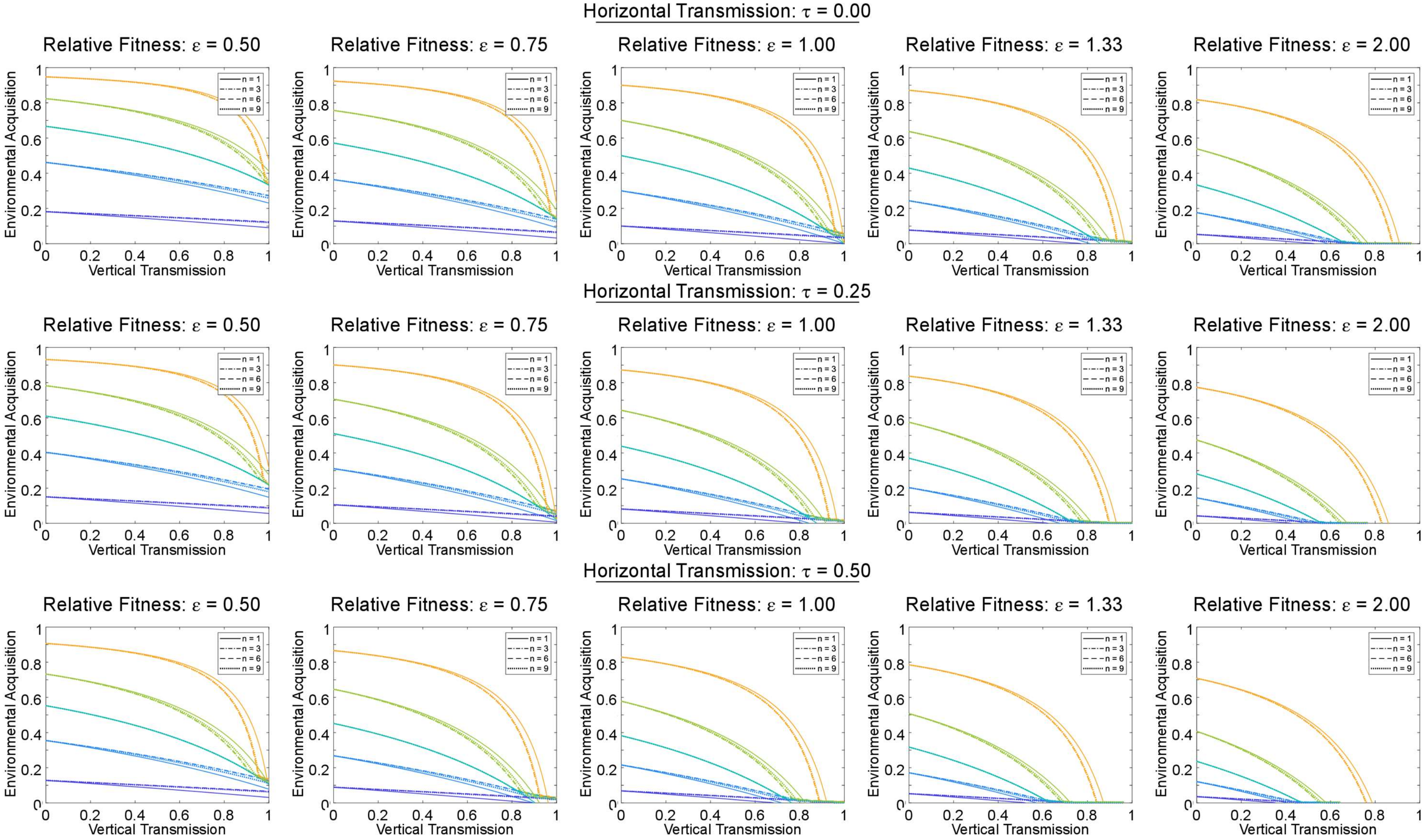


**Fig S3A. Effect of transmission modes, host mate choice, and microbe effects on host fitness on host-microbe carrier frequencies within a single population of sexually reproducing hosts. Females are the choosing sex and prefer males of the same carrier/non-carrier status.** Here each line represents different steady-state microbe carrier frequencies (0.1-0.9, increasing in 0.2 intervals). From top to bottom: increasing levels of horizontal transmission for 0-0.5. From left to right: relative fitness of carriers to non-carriers 0.5-2. Here the effects of mate choice on carrier frequencies are less pronounced than for male mate choice compare with Fig S5.

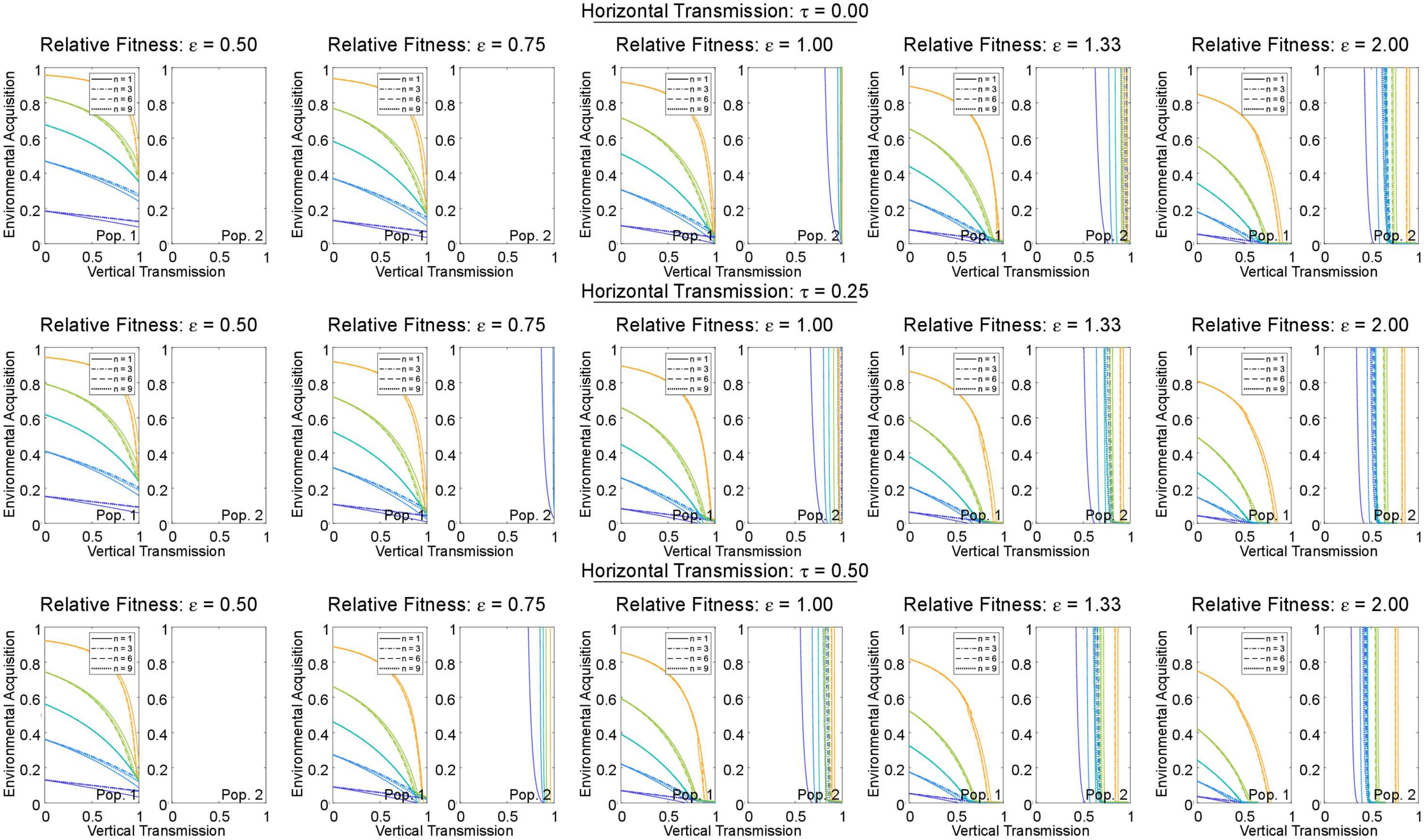


**Fig S3B. Effect of transmission modes, host mate choice, and microbe effects on host fitness on host-microbe carrier frequencies within two populations of sexually reproducing hosts. Females are the choosing sex and prefer males of the same carrier/non-carrier status.** Here each line represents different steady-state microbe carrier frequencies (0.1-0.9, increasing in 0.2 intervals). From top to bottom: increasing levels of horizontal transmission for 0-0.5. From left to right: relative fitness of carriers to non-carriers 0.5-2. In population 1 microbes can proliferate independently of hosts and there is environmental acquisition by the host. In population 2 microbes cannot be acquired by hosts from the environment, only via maternal VT or horizontal transmission from migrants from population 1.

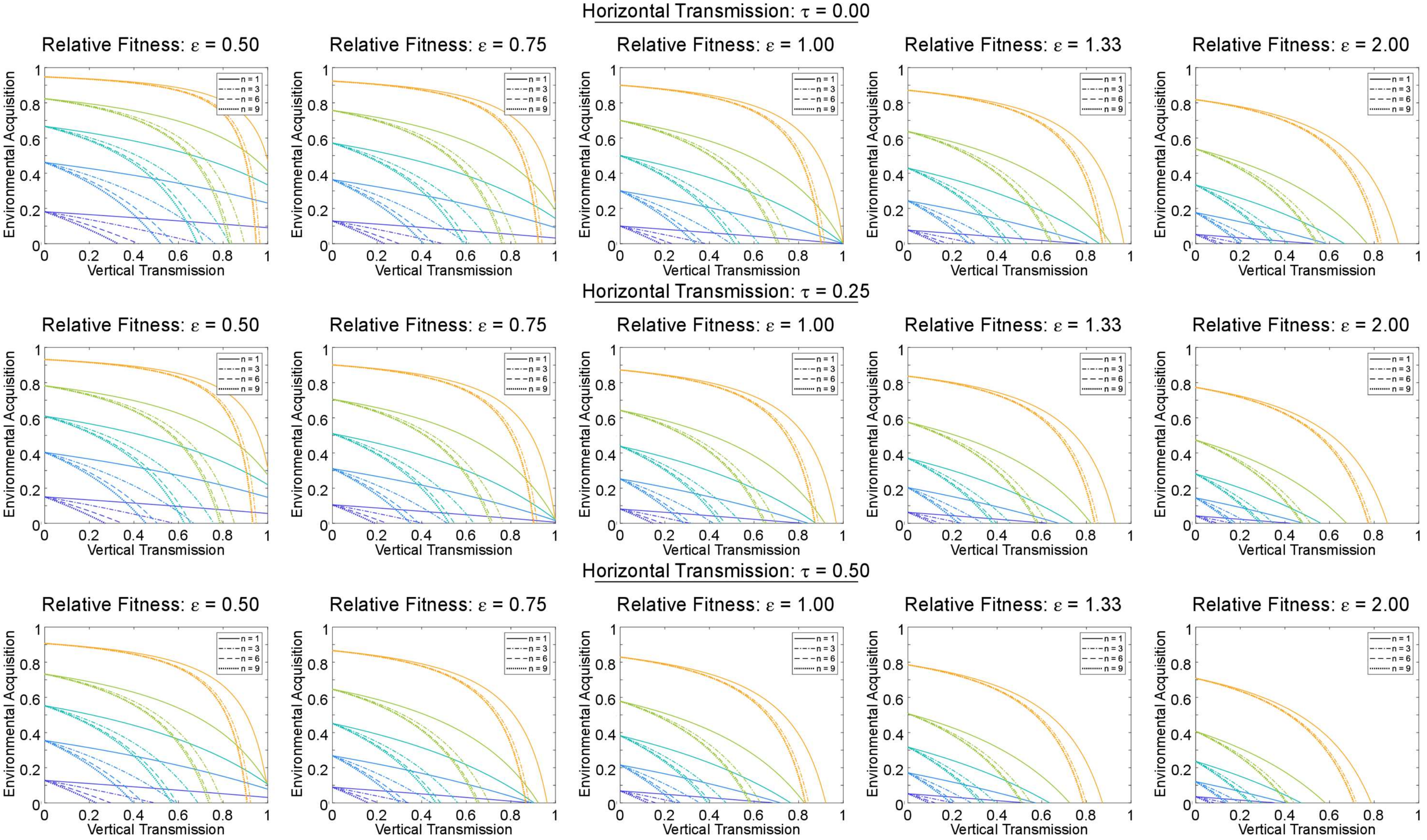


**Fig S4A. Effect of transmission modes, host mate choice, and microbe effects on host fitness, on host-microbe carrier frequencies within a single population of sexually reproducing hosts. Males are the choosing sex and prefer females that are microbe-carriers.** Here each line represents different steady-state microbe carrier frequencies (0.1-0.9, increasing in 0.2 intervals). From top to bottom: increasing levels of horizontal transmission for 0-0.5. From left to right: relative fitness of carriers to non-carriers 0.5-2. Top row panels 2 & 4 (Left to Right) are reproduced in main text Fig 2C and D.

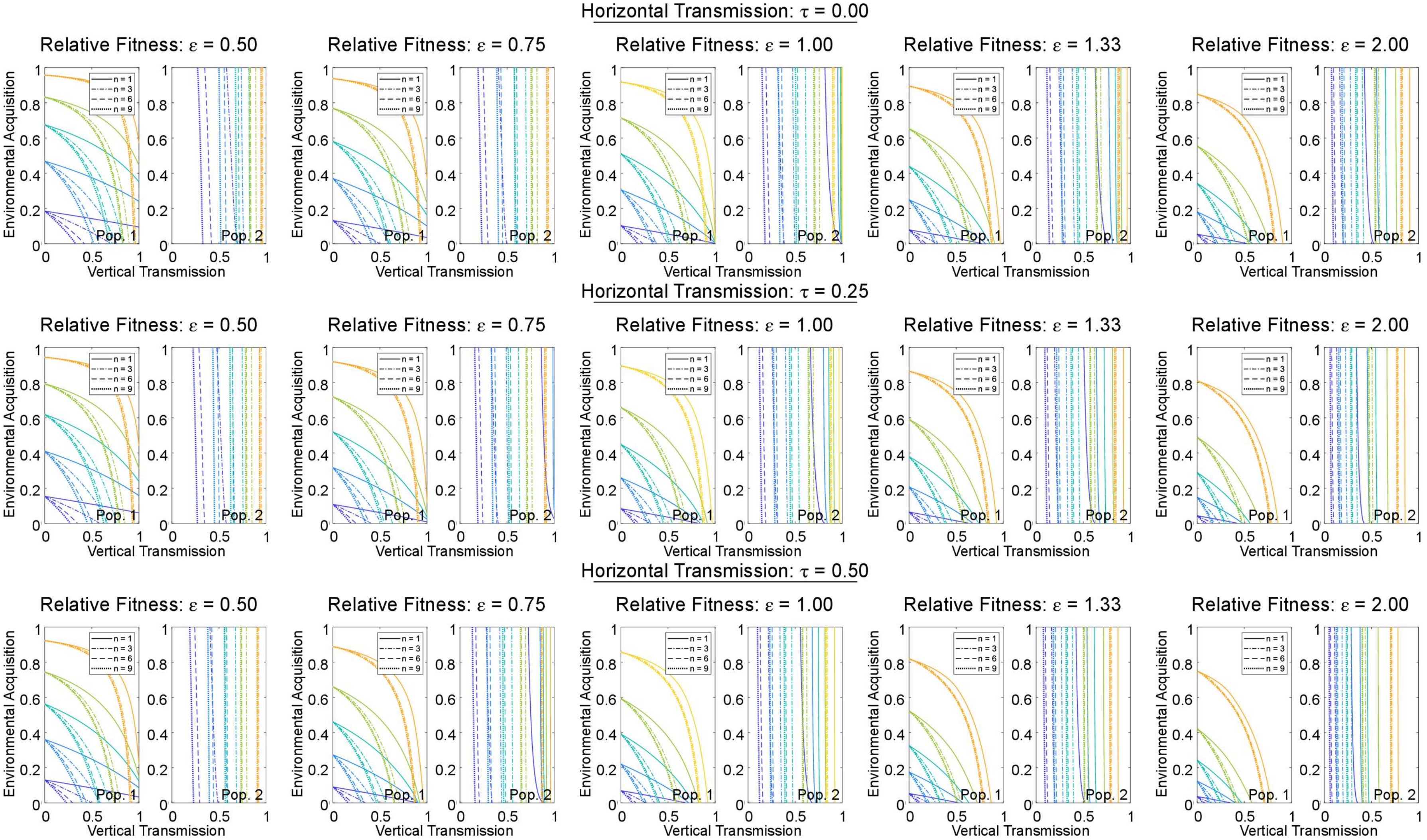


**Fig S4B. Effect of transmission modes, host mate choice, and microbe effects on host fitness on host-microbe carrier frequencies within two populations of sexually reproducing hosts. Males are the choosing sex and prefer females that are microbe-carriers.** Here each line represents different steady-state microbe carrier frequencies (0.1-0.9, increasing in 0.2 intervals). From top to bottom: increasing levels of horizontal transmission for 0-0.5. From left to right: relative fitness of carriers to non-carriers 0.5-2. In population 1 microbes can proliferate independently of hosts and there is environmental acquisition by the host. In population 2 microbes cannot be acquired by hosts from the environment, only via maternal VT or horizontal transmission from migrants from population 1. Top row panels 3 & 5 (Left to Right) are reproduced in main text Fig 3C and D.

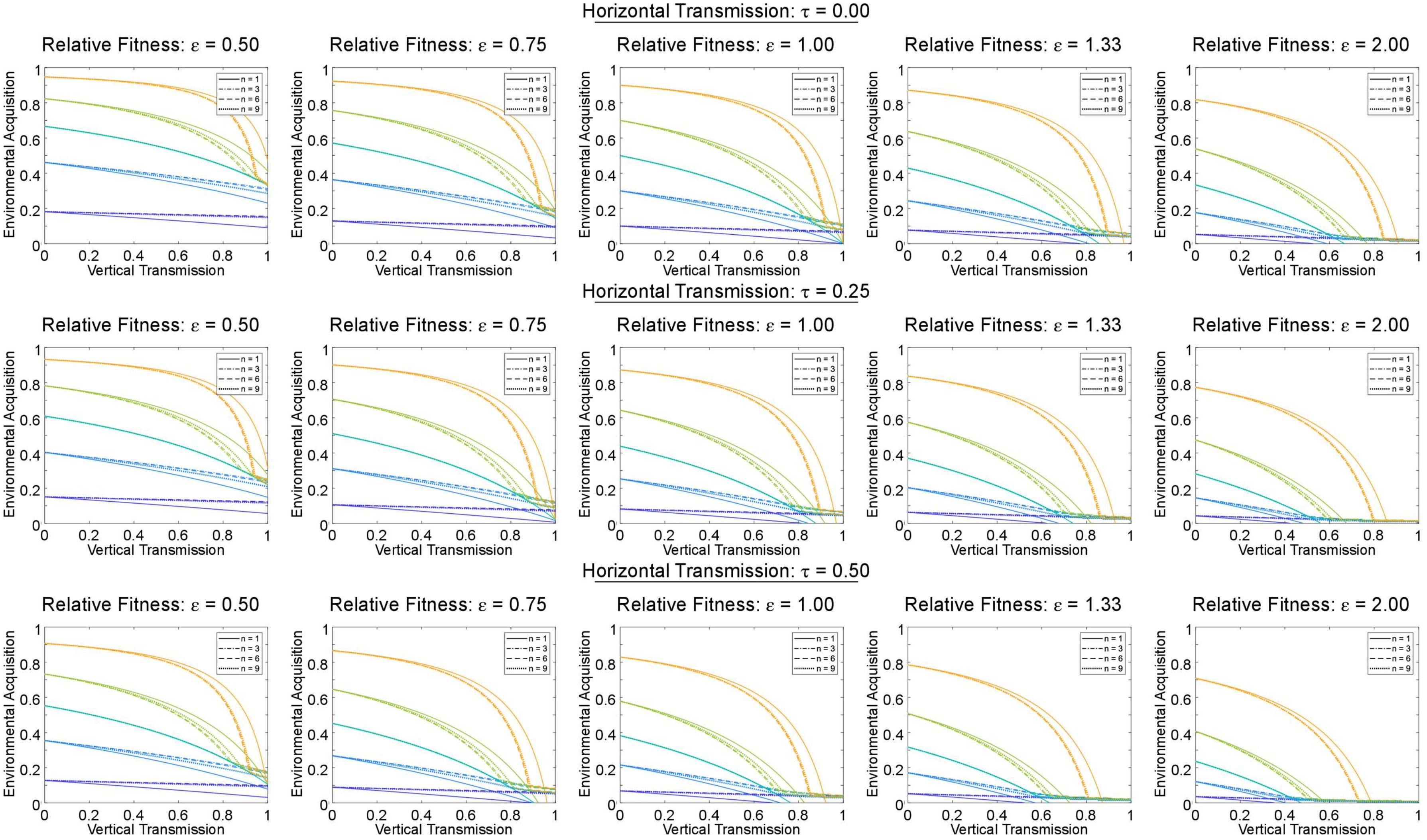


**Fig S5A. Effect of transmission modes, host mate choice, and microbe effects on host fitness on host-microbe carrier frequencies within a single population of sexually reproducing hosts. Males are the choosing sex and prefer females of the same carrier/non-carrier status.** Here each line represents different steady-state microbe carrier frequencies (0.1-0.9, increasing in 0.2 intervals). From top to bottom: increasing levels of horizontal transmission for 0-0.5. From left to right: relative fitness of carriers to non-carriers 0.5-2. Top row panels 2 & 4 (Left to Right) are reproduced in main text Fig 2A and B.

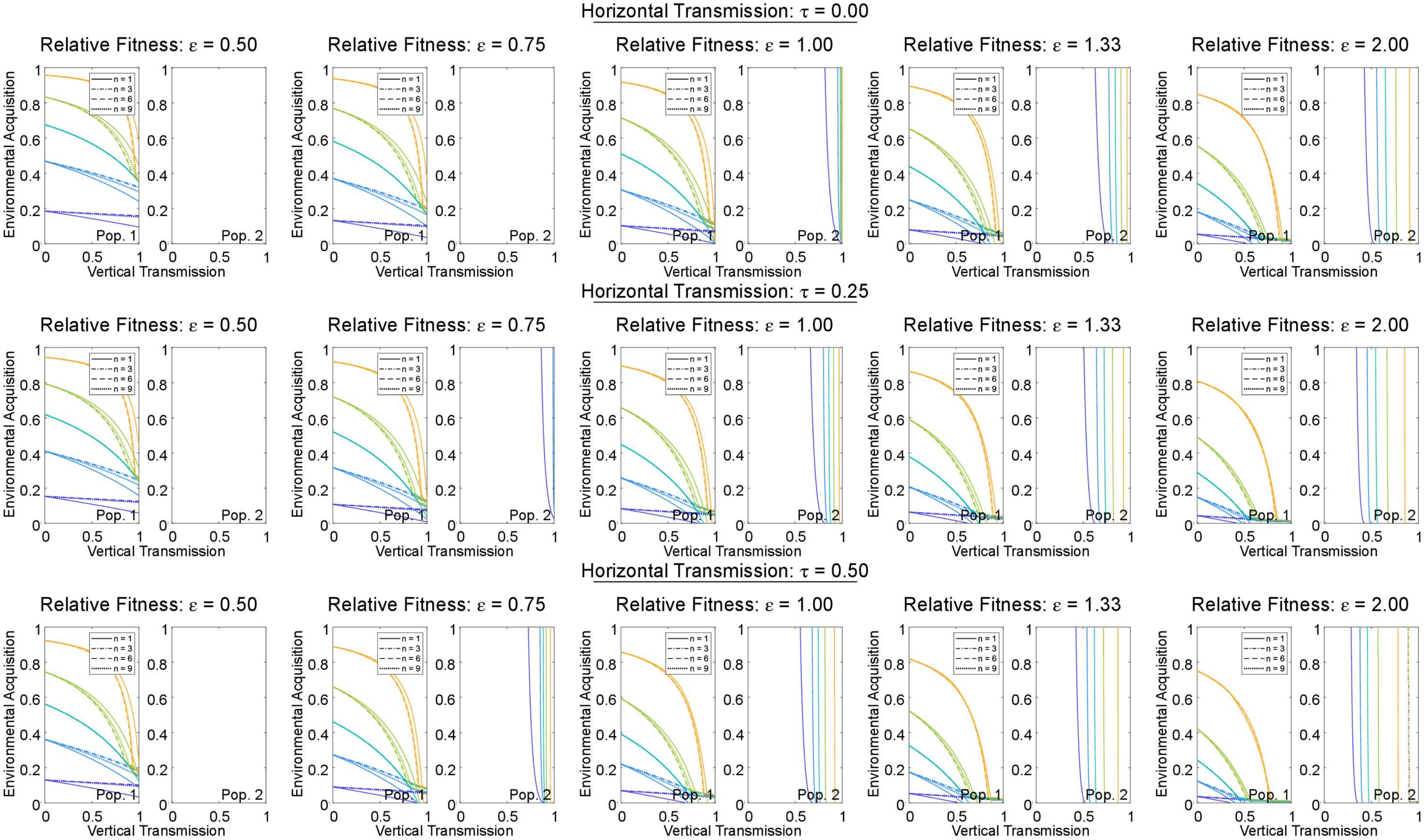


**Fig S5B. Effect of transmission modes, host mate choice, and microbe effects on host fitness on host-microbe carrier frequencies within two populations of sexually reproducing hosts. Males are the choosing sex and prefer females of the same carrier/non-carrier status.** Here each line represents different steady-state microbe carrier frequencies (0.1-0.9, increasing in 0.2 intervals). From top to bottom: increasing levels of horizontal transmission for 0-0.5. From left to right: relative fitness of carriers to non-carriers 0.5-2. In population 1 microbes can proliferate independently of hosts and there is environmental acquisition by the host. In population 2 microbes cannot be acquired by hosts from the environment, only via maternal VT or horizontal transmission from migrants from population 1. Top row panels 3 & 5 (Left to Right) are reproduced in main text Fig 3A and B.
